# Efficacy of Guanabenz Combination Therapy Against Chronic Toxoplasmosis Across Multiple Mouse Strains

**DOI:** 10.1101/2020.03.19.999672

**Authors:** Jennifer Martynowicz, J. Stone Doggett, William J. Sullivan

## Abstract

*Toxoplasma gondii*, an obligate intracellular parasite that can cause life-threatening acute disease, differentiates into a quiescent cyst stage to establish lifelong chronic infections in animal hosts, including humans. This tissue cyst reservoir, which can reactivate into an acute infection, is currently refractory to clinically available therapeutics. Recently, we and others have discovered drugs capable of significantly reducing brain cyst burden in latently infected mice, but not to undetectable levels. In this study, we examined the use of novel combination therapies possessing multiple mechanisms of action in mouse models of latent toxoplasmosis. Our drug regimens included combinations of pyrimethamine, clindamycin, guanabenz, and endochin-like quinolones (ELQs), and were administered to two different mouse strains in an attempt to eradicate brain tissue cysts. We observed mouse strain-dependent effects with these drug treatments: pyrimethamine + guanabenz showed synergistic efficacy in C57BL/6 mice, yet did not improve upon guanabenz monotherapy in BALB/c mice. Contrary to promising *in vitro* results demonstrating toxicity to bradyzoites, we observed an antagonistic effect between guanabenz + ELQ-334 *in vivo*. While we were unable to completely eliminate brain cyst burden, we found that a combination treatment of ELQ-334 + pyrimethamine impressively reduced brain cysts to 95% in C57BL/6 mice, which approaches the limit of detection. These analyses highlight the importance of evaluating anti-infective drugs in multiple mouse strains and will help inform further preclinical cocktail therapy studies designed to treat chronic toxoplasmosis.

## Introduction

*Toxoplasma gondii* is an obligate intracellular parasite of warm-blooded vertebrates, and it has infected up to one-third of humans (1). The parasite can be transmitted across three life cycle stages: proliferative tachyzoites can cross the placenta and infect the fetus, latent bradyzoites can be consumed in raw or undercooked meat, and oocysts excreted by the definitive host (felines) can be ingested or inhaled (2). The destructive lytic nature of tachyzoites can cause widespread damage throughout the host and is lethal without treatment (3). However, in an immune competent host, tachyzoites rapidly convert into bradyzoites, which reside in intracellular tissue cysts that form in muscle and organs, including the brain (4-6). The bradyzoite-containing tissue cysts are responsible for long-lasting chronic infection.

Upon immunosuppression of the host, bradyzoites convert into the lytic tachyzoites and cause acute disease. Individuals who are at risk for reactivation of acute infection are placed on prophylaxis until they can return to immune competency. For hematopoietic cell transplant patients and solid organ transplant patients who develop toxoplasmosis, lifelong secondary prophylaxis may be required (7). The current frontline treatment of antifolates (pyrimethamine + sulfadiazine) suffers serious shortcomings (8). Up to a third of patients are allergic to sulfa drugs and there is a high rate of adverse effects to pyrimethamine, most notably hematologic toxicity. These adverse events are noted in up to 60% of *Toxoplasma* encephalitis patients, which leads to discontinuation of therapy in up to 45% of patients (3, 9, 10). Additional side effects include leukopenia, thrombocytopenia, cutaneous rash, and fever (10). Most importantly, no approved treatment effectively targets tissue cysts. It is therefore of clinical importance to identify better tolerated agents that can eliminate the tissue cyst reservoir that constantly jeopardizes the life of susceptible individuals.

Targeting tissue cysts remains a technical challenge due to their quiescent nature and proclivity for the brain. Only in the last decade have a few compounds capable of significantly reducing tissue cysts *in vivo* appeared. Compound 32, which was optimized for use against *T. gondii* calcium-dependent protein kinase 1 (TgCDPK1), shows an 88.7% decrease in brain cyst burden in BALB/c mice (11). Endochin-like quinolones (ELQs) that target the parasite cytochrome *bc*_1_ complex have also shown promise in lowering cyst counts in mouse models (12, 13). ELQ-316 in particular reduces brain cyst burden 76-88% in CBA/J mice (14).

Previous work in our lab found that the FDA-approved anti-hypertensive drug, guanabenz, can be repurposed as an anti-parasitic (15-17). As reported in other eukaryotic cells, guanabenz can inhibit the dephosphorylation of translation initiation factor eIF2α in *Toxoplasma*, which is a key switch associated with stage conversion (15). We have shown that guanabenz inhibits growth of tachyzoites and reduces brain cyst burden in chronically infected BALB/c mice 65-80% (16). Interestingly, guanabenz had a detrimental effect on pathogenesis in C57BL/6 mice, which are genetically more susceptible to *Toxoplasma* infection (17). Half of the chronically infected mice treated with guanabenz group died and the cyst burden increased in the surviving C57BL/6 mice. The efficacy of compound 32 and ELQs in the more susceptible mouse strain C57BL/6 has not been determined.

A common shortfall of these three cyst-reducing compounds is that none achieve a radical cure—for unknown reasons, they are unable to completely eliminate *Toxoplasma* from the brain. Eradicating this residual cyst population is not a simple matter of prolonging treatment time. We have previously attempted prolonging guanabenz treatment to 6 weeks but were unable to improve cyst burden reduction (17). Additionally, a residual population of cysts was still maintained after prolonged treatment with ELQs (*Doggett, jointly submitted manuscript*).

While the reduction is brain cysts with these novel compounds is promising, the ideal treatment needs to eliminate them completely. In this study, we tested the hypothesis that combination treatments coupling guanabenz with various drugs possessing different mechanisms of action would lower cyst burden beyond monotherapy.

## Methods

### Parasite strains and culture

*Toxoplasma gondii* parasites (Type II Prugniad (Pru) strain) were collected as tissue cysts from brains of chronically infected BALB/c mice and used to infect human foreskin fibroblasts (HFF). The infected cultures were maintained in Dulbecco’s medium supplemented with 1% heat-inactivated fetal bovine serum (FBS) in a humidified incubator at 37°C with 5% CO_2_. To ensure their developmental capacity, tachyzoites were maintained in culture no more than 15 passages. Parasites were tested for mycoplasma using PCR as previously described (18) prior to use in mice.

### Mouse strains and infection

The mice used in this study were housed in American Association for Accreditation of Laboratory Animal Care (AAALAC) approved facilities at either the Indiana University School of Medicine Laboratory Animal Research Center (LARC) or the IUPUI Science Animal Research Center (SARC). The Institutional Animal Care and Use Committee (IACUC) at Indiana University School of Medicine approved the use of all animals and procedures (IACUC protocol number 11376).

BALB/c mice were purchased from the Jackson Laboratory (Bar Harbor, ME) at 5 weeks old. The mice were allowed to acclimate for one week before being intraperitoneally (i.p.) infected with 10^4^ Pru tachyzoites suspended in 100μl of autoclaved, filter-sterilized PBS. C57BL/6 mice, also purchased from the Jackson Laboratory at 5 weeks of age, were infected with 10^3^ Pru tachyzoites. Mice were routinely observed multiple times a day throughout the course of acute infection. After 21 days post-infection, mice were randomized into treatment groups. Blood samples were extracted by cardiac puncture to confirm parasite infection through serological analysis of collected serum to identify parasite specific antibodies using a dot blot containing parasite lysate.

### Drug sources and administration

Stock solutions were made at the beginning of the experiment and aliquoted in 1.5mL Eppendorf tubes and stored at 4°C until needed. Guanabenz (Sigma) was delivered i.p. 5mg/kg/day in sterile saline. Pyrimethamine (MP Biomedicals) and clindamycin (Sigma) were dosed i.p. at 20mg/kg/day in sterile saline. ELQ-316 and ELQ-334 were synthesized as described in the accompanying publication (*Doggett, jointly submitted manuscript*), identified by proton nuclear magnetic resonance (^1^H-NMR), and determined to be >95% pure by reversed-phase high-performance liquid chromatography (HPLC). ELQ-334 was administered by oral gavage using a flexible plastic gavage needle at 10mg/kg/day in corn oil. Treatment time for each drug was three weeks.

### Brain Cyst Quantification

Brains were extracted upon euthanasia and cyst burden was quantified as previously described (16). Briefly, the dissected tissue was homogenized in 650μl of sterile PBS using a mortar and pestle. A 250ul aliquot of homogenate was fixed using 3% methanol-free formaldehyde for 20 minutes. The homogenate was blocked using 3% bovine serum albumin (BSA) in 0.2% Triton X-100 before staining with 1:250 rhodamine-conjugated *Dolichos biflorus* lectin (Vector Laboratories Inc.) to visualize the cyst wall. Five microliter aliquots of stained homogenate were placed on a coverslip and sealed before imaging. Stained samples were blinded before cysts were counted under 20x magnification. The counted value was then extrapolated to estimate the cyst burden for the entire brain.

### *In vitro* parasite viability assays

For the plaque assay, 12-well plates containing confluent HFF monolayers were infected with 10^4^ Pru tachyzoites and allowed to invade for 2 hours. The plates were then washed with warm sterile PBS twice before fresh medium supplemented with the indicated drug was added. Plates were left undisturbed under normal culture conditions for 13 days. They were then fixed with 100% ice cold methanol for 20 minutes and stained with crystal violet for 20 minutes. For parasite doubling assays, 12-well plates of HFFs were infected with 10^3^ Pru tachyzoites and allowed to invade for 2 hours. The plates were then washed with warm sterile PBS twice before fresh medium supplemented with the indicated drug was added. Plates were left undisturbed under normal culture conditions for 36 hours. They were then fixed with 100% ice cold methanol for 20 minutes and washed with PBS. The number of parasites within 50 randomly chosen vacuoles was counted.

### Statistics

Statistical analysis was performed using GraphPad Prism version 7.03. For datasets comparing two groups, statistical significance was determined using Student’s t test. If more than two groups were compared, the datasets were analyzed using One Way AVOVA to determine statistical significance before secondary analysis with Student’s t test. Survival data was analyzed using a log rank test. P values of ≤0.05 were considered statistically significant.

## Results

### Pyrimethamine prevents guanabenz-associated death in C57BL/6 mice and reduces brain cyst burden

We previously showed that while guanabenz effectively reduces brain cyst burden in BALB/c mice, it increases cyst burden in C57BL/6 mice (17). The C57BL/6 mice also developed symptoms of *Toxoplasma* encephalitis during guanabenz treatment, resulting in loss of half the treatment group. We postulated that the guanabenz treatment led to reactivation of latently encysted parasites in C57BL/6 mice. In an attempt to prevent the death of chronically infected C57BL/6 mice treated with guanabenz, we co-administered pyrimethamine; pyrimethamine is the frontline antifolate used to treat acute toxoplasmosis by specifically targeting replicating parasites (Fig. 1A). As seen previously (17), treatment of latently infected C57BL/6 mice with guanabenz alone resulted in death of half the group (Fig. 1B). In contrast, none of the infected C57BL/6 mice co-administered guanabenz and pyrimethamine perished (Fig. 1B). The efficacy of pyrimethamine with guanabenz suggests that the detrimental effects of guanabenz treatment in C57BL/6 mice can be nullified with combination therapy.

**Figure 1.**
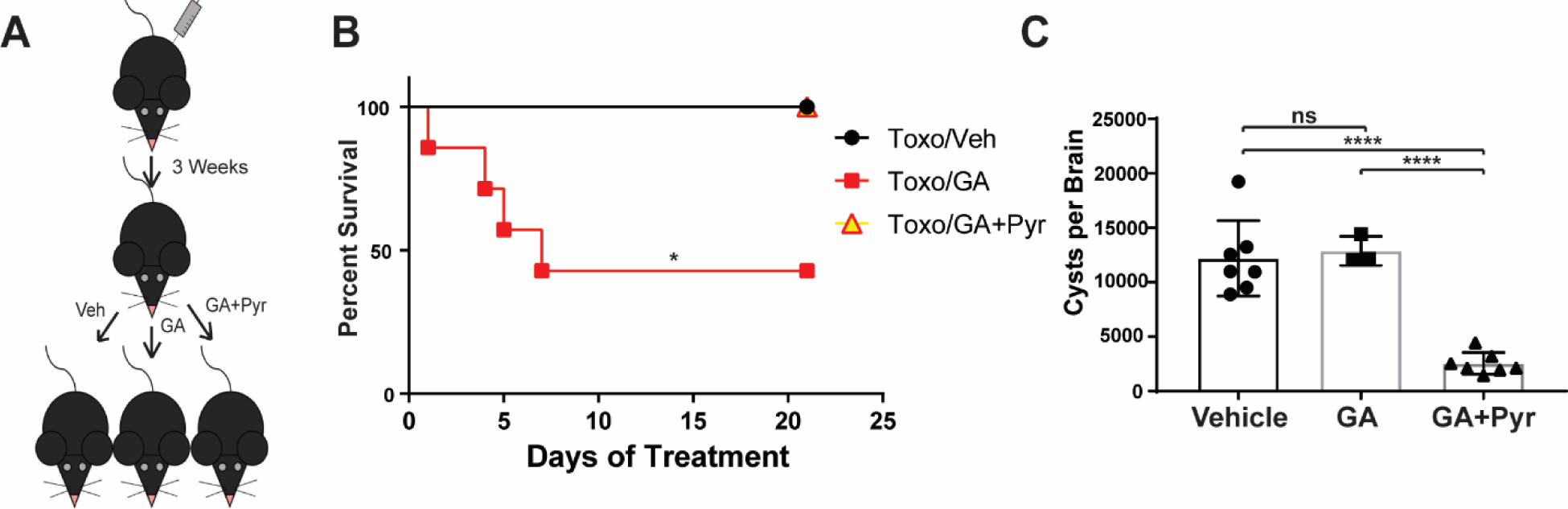
Pyrimethamine combination therapy in C57BL/6 mice. (A) Six-week old, female C57BL/6 mice were infected intraperitoneally with *Toxoplasma* (grey syringe) and allowed to progress to chronic infection for three weeks. The mice were then randomized into three different treatment groups. N=7 (B) The latently infected mice were treated for three weeks and survival was tracked over that time. Log rank test, *p= 0.0221. (C) At the conclusion, brain tissue cysts burden was blindly quantified for each mouse. One way ANOVA p=<0.0001 followed by an unpaired Student’s t test. ns=not significant, ****p<0.0001.

At the conclusion of the study, brains were collected and cyst burden was blindly quantified. As seen before (17), guanabenz alone failed to reduce cyst counts in the brains of C57BL/6 mice (Fig. 1C). Surprisingly, combination therapy with guanabenz and pyrimethamine restored the cyst-lowering capabilities of guanabenz in C57BL/6 mice (Fig. 1C), comparable to levels observed in BALB/c mice with chronic toxoplasmosis treated with guanabenz (16, 17). These findings show that coupling guanabenz with a drug that potently targets tachyzoites is more efficacious in reducing cyst counts in latently infected mice.

### Co-administration of pyrimethamine or clindamycin with guanabenz in BALB/c mice

In contrast to C57BL/6 mice, which are more susceptible to *Toxoplasma* infection, guanabenz alone is sufficient to significantly lower brain cyst counts in latently infected BALB/c mice (16, 17). Given the synergy of the combination treatment in C57BL/6 mice, we sought to apply it to the BALB/c model of infection in hopes that it would completely eliminate cysts in these more resistant mice. Moreover, in addition to pyrimethamine, we tested a combination of guanabenz with another clinically used anti-parasitic effective against tachyzoites: clindamycin (Fig. 2A) (19-21). Unfortunately, neither the addition of pyrimethamine or clindamycin with guanabenz showed further reduction in brain cysts than guanabenz alone after 3 weeks of treatment (Fig. 2B). These findings indicate that a residual population of brain cysts (∼20-30%) remains in BALB/c mice when guanabenz is administered alone or in combination with an anti-tachyzoite agent.

**Figure 2.**
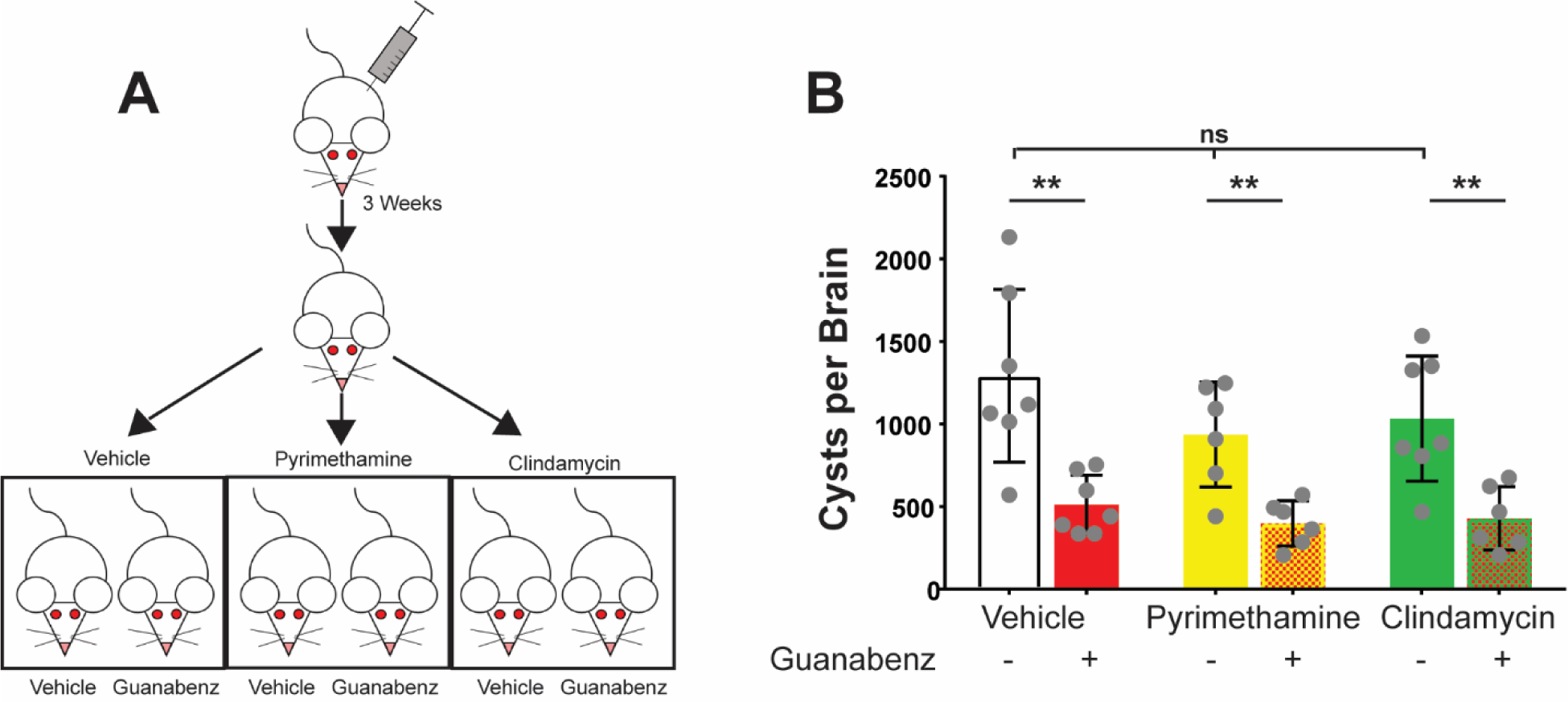
Effect of pyrimethamine or clindamycin with guanabenz on BALB/c brain cyst burden. (A) Six-week old, female BALB/c mice were infected intraperitoneally with *Toxoplasma* (grey syringe) and allowed to progress to chronic infection for three weeks. The latently infected mice were then randomized into six different treatment groups. N=7 (B) Following three weeks of indicated drug treatment, brain cyst burden was blindly quantified. One way ANOVA p<0.0001followed by an unpaired Student’s t test. ns=not significant, **p<0.01.

### Effects of guanabenz combined with ELQ against *Toxoplasma*

The current therapeutics against toxoplasmosis fail to eradicate the infection because they do not have significant activity against tissue cysts. We hypothesized that combination therapies composed of drugs with cyst-reducing capabilities may have synergistic effects. Guanabenz and endochin-like quinolones (ELQs) exhibit activity against tachyzoites and reduce brain cyst counts 70-80% through different mechanisms of action (12, 14). To address our hypothesis, we first examined the effects of guanabenz, ELQ-316, or both on Pru strain tachyzoites in culture. A plaque assay showed an additive effect for guanabenz and ELQ-316, with minimal plaques at low doses and no visible plaques at the high doses (Fig. 3A). A synergistic activity against tachyzoite replication was also observed in a parasite doubling assay (Fig. 3B). The same synergistic activity against tachyzoites was observed for another type II parasite strain, ME49 (data not shown).

**Figure 3.**
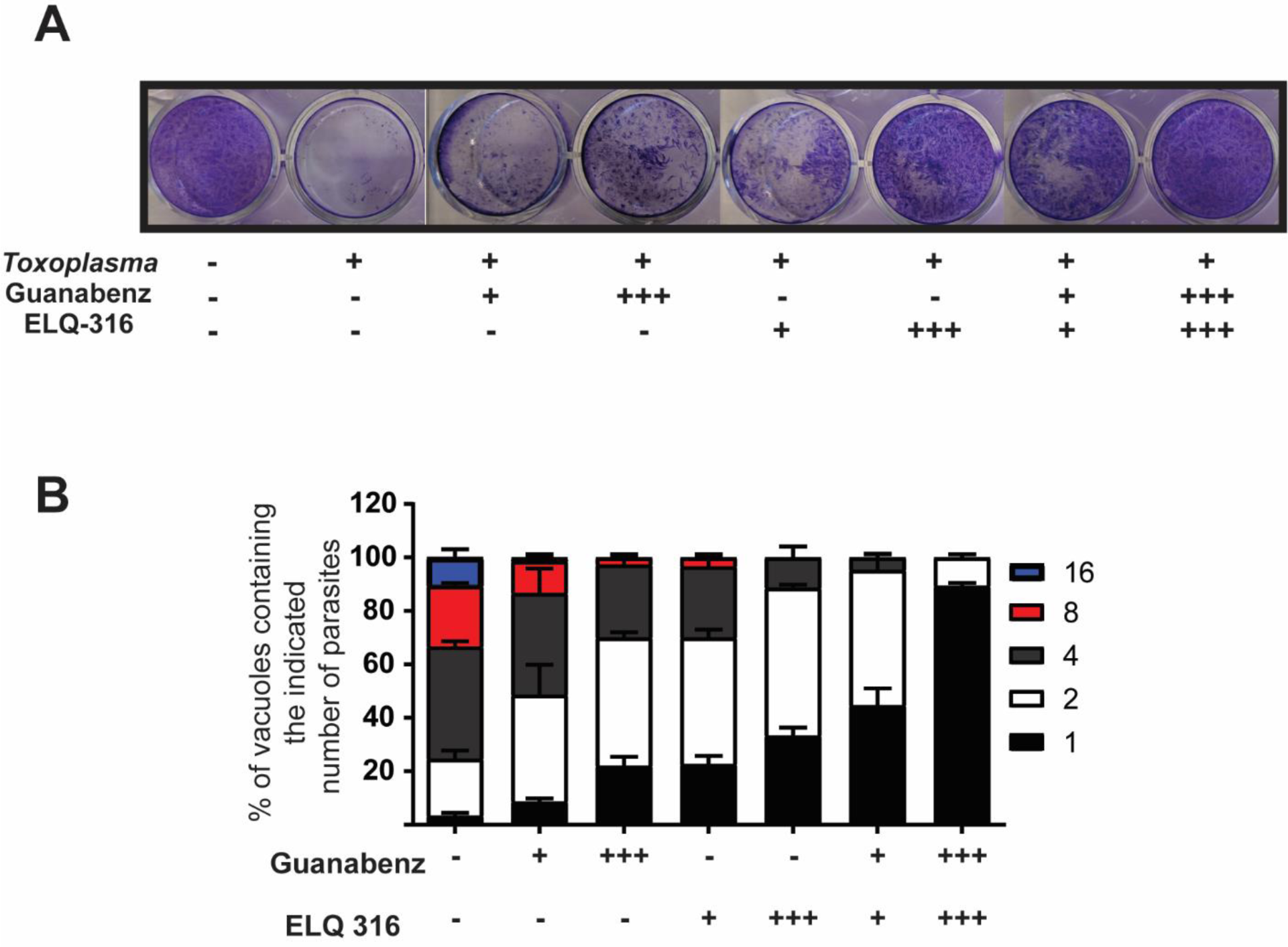
Additive effects of guanabenz and ELQ-316 in vitro. (A) Plaque assay of an HFF monolayer with or without *Toxoplasma* and the indicated drug concentrations. (B) Doubling assay of Pru tachyzoites after 36 hours of drug treatment, performed with three biological replicates done in technical triplicate. For both A and B, (+) denotes IC_50_ of drug and (+++) denotes 3x IC_50_.

When guanabenz is added to tissue cysts generated *in vitro*, the bradyzoites take on an anomalous morphology. Bradyzoites within the guanabenz-treated cysts appear to shrivel up, resulting in large empty spaces within the cyst (Fig. 4, black arrow). Treatment of tissue cysts with ELQ-316 was also damaging, resulting in cysts that appear shrunken and collapsed (Fig. 4, yellow arrow). In addition to using *Dolichos* lectin to stain cyst walls, we stained these cultures for the tachyzoite surface antigen SAG1. No significant staining with SAG1 was evident in tissue cyst cultures treated with guanabenz alone. However, in the ELQ-316 treated tissue cysts, an increased number of SAG1 positive parasites was observed (Fig. 4, grey arrow). When the two drugs are administered together, SAG1 positive parasites were evident both within and outside of tissue cysts, suggesting that these treated bradyzoites may be converting to tachyzoites before dying.

**Figure 4.**
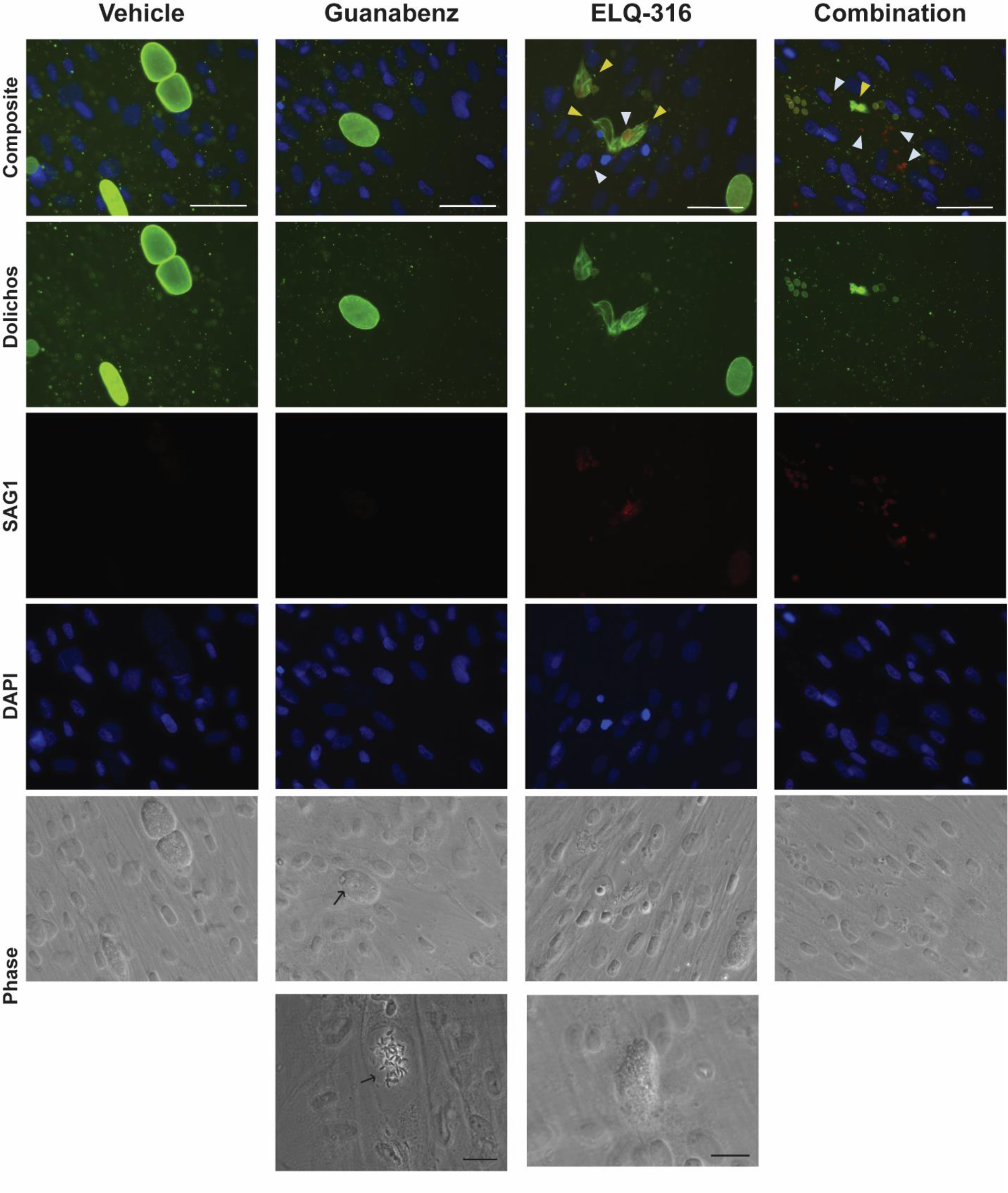
Guanabenz and ELQ-316 cause abnormal cyst morphology. Bradyzoite cysts were generated by pH stress for 5 days in vitro before being drug-treated for four days. Black arrow denotes swollen cyst structure with disorganized parasites inside. Yellow arrowhead denotes shrunken cyst wall. Grey arrowhead = SAG1+ parasites. White scale bar=50μM; black scale bar=20μM.

Given the promising results *in vitro*, we proceeded to test the efficacy of guanabenz and ELQ combination therapy in chronically infected BALB/c mice (Fig. 5A). ELQ-334 is a prodrug form of ELQ-316 designed to have increased bioavailability and stability (22). BALB/c mice with latent toxoplasmosis were treated with guanabenz, ELQ-334, or both for 3 weeks. As expected, both guanabenz and ELQ-334 monotherapy reduced brain cysts; surprisingly, the combination therapy had no effect on cyst counts (Fig. 5B). Rather than being synergistic, these drugs nullify one another’s ability to reduce brain cysts *in vivo*.

**Figure 5.**
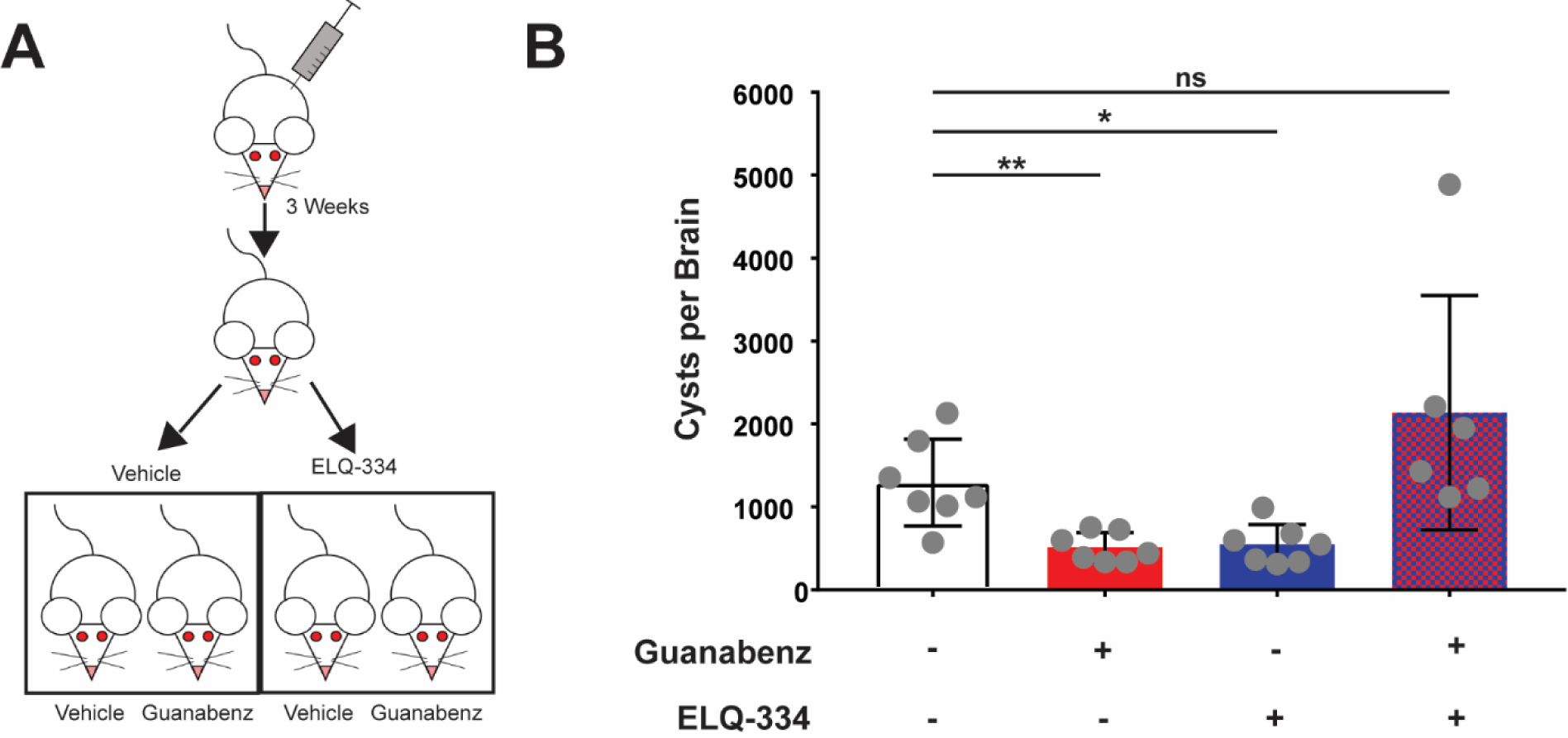
ELQ-334 does not have an additive effect with guanabenz in BALB/c mice. (A) Six-week old, female BALB/c mice were infected intraperitoneally with *Toxoplasma* (grey syringe) and allowed to progress to chronic infection for three weeks. The latently infected mice were then randomized into four different treatment groups. N=7 (B) Following three weeks of indicated drug treatment, brain cyst burden was blindly quantified. One way ANOVA p=0.0015 followed by an unpaired Student’s t test. ns=not significant, *p<0.05, **p<0.01.

### Guanabenz+ELQ+pyrimethamine does not outperform monotherapy

Figure 4 reveals a possible explanation for the failed guanabenz+ELQ combination therapy: the SAG1+ parasites observed in drug-treated cyst cultures could be a source of tachyzoites that reseed the brain with cysts. To test this possibility, we added pyrimethamine to the guanabenz+ELQ therapy (Fig. 6A).

**Figure 6.**
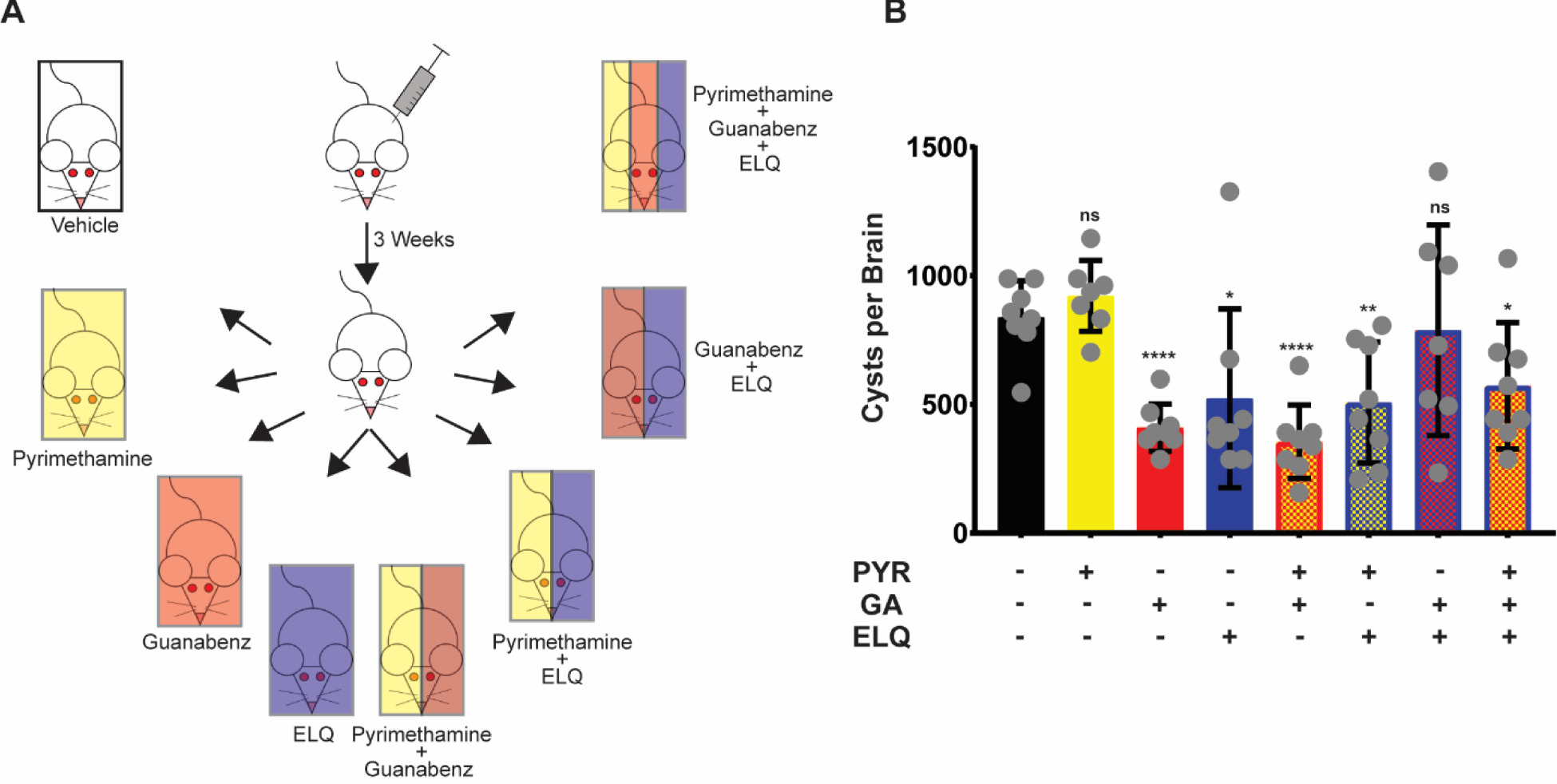
Triple combination therapy in BALB/c mice. (A) Six-week old, female BALB/c mice were infected intraperitoneally with *Toxoplasma* (grey syringe) and allowed to progress to chronic infection for three weeks, then randomized into eight different treatment groups. (B) Following three weeks of indicated drug treatment, brain cyst burden was blindly quantified. One way ANOVA p<0.0001 followed by an unpaired Student’s t test. Statistics shown are compared to vehicle control. ns=not significant, *p<0.05, **p<0.01, ****p<0.0001.

The experimental set up accounted for all possible combinations of the three drugs in an attempt to better understand the mechanism for the failure of the guanabenz+ELQ treatment.We again observed no reduction in brain cysts when guanabenz and ELQ-334 were co-administered, but were able to reestablish a statistically significant reduction in cyst burden when all three drugs were given (Fig. 6B). However, the triple therapy did not significantly lower the cyst burden below that of the guanabenz or ELQ monotherapies.

Since C57BL/6 mice are more sensitive to tachyzoites, we sought to confirm the improved efficacy of the triple therapy in this mouse strain. We hypothesized that the triple therapy would display better efficacy in C57BL/6 mice based on the improvement seen when pyrimethamine was combined with guanabenz (Fig. 1C). A schematic of the experimental design is shown in Figure 7A.

**Figure 7.**
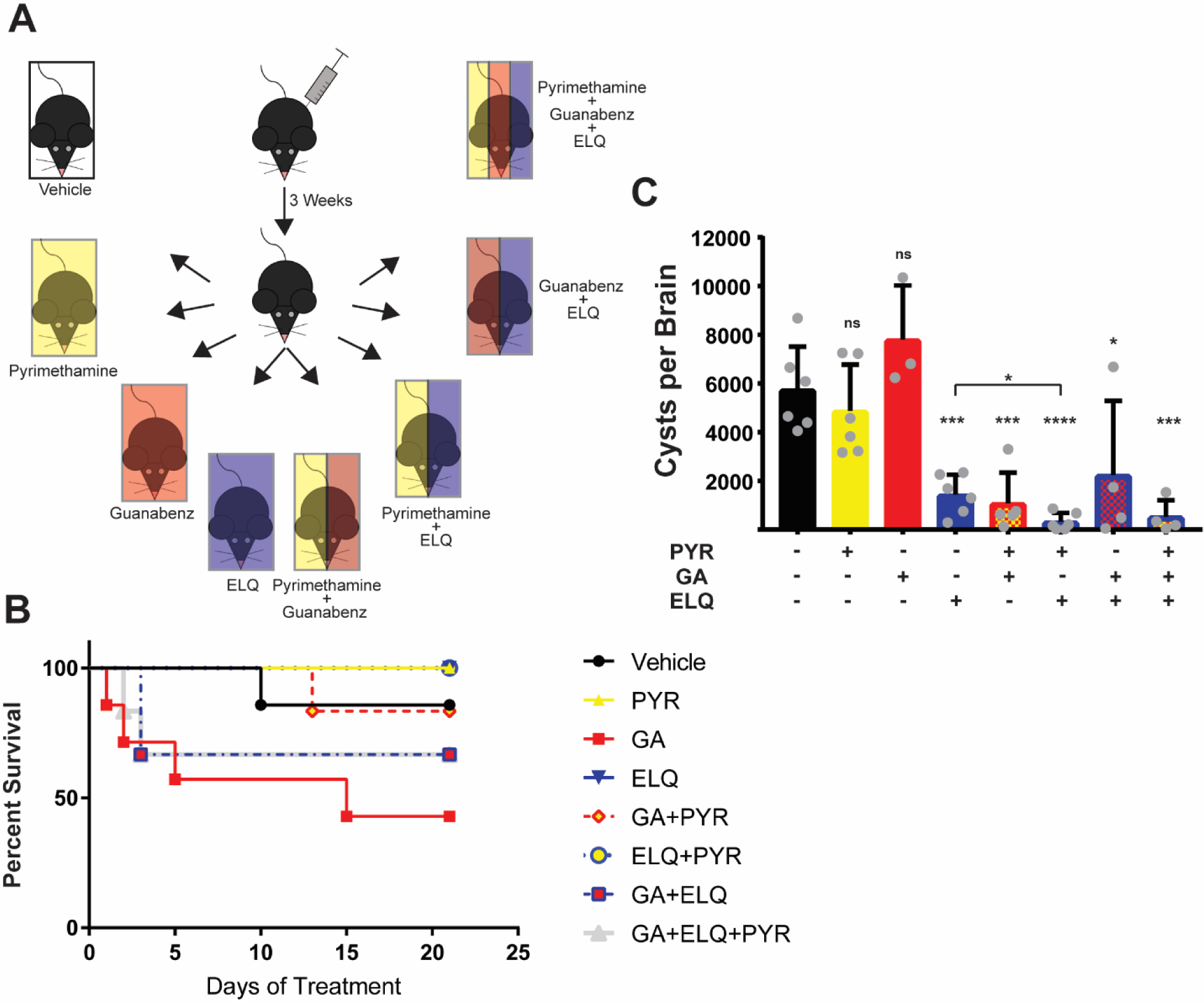
Triple combination therapy in C57BL/6 mice. (A) Six-week old, female C57BL/6 mice were infected intraperitoneally with *Toxoplasma* (grey syringe) and allowed to progress to chronic infection for three weeks, then randomized into eight different treatment groups. N=6 (B) The latently infected mice were treated with indicated drugs for three weeks and survival was tracked over that time. Not statistically significant by log-rank test. (C) Following three weeks of treatment, brain cyst burden was blindly quantified. One way ANOVA p<0.0001 followed by an unpaired Student’s t test. Statistics above the bar are compared to vehicle control unless otherwise shown. ns=not significant, *p<0.05, **p<0.01, p<0.001, ****p<0.0001.

Given that the combination of guanabenz and ELQ-334 failed to reduce brain cysts in BALB/c mice (Fig. 5B), we were not surprised to see reduced survival in some of the latently infected C57BL/6 mice treated with this combination, though it did not reach statistical significance (Fig. 7B). However, the inclusion of pyrimethamine did not improve survival rates, which suggests continued failure of the triple therapy in controlling infection (Fig. 7B). Brain cyst burdens in the survivors of the guanabenz+ELQ and the guanabenz+ELQ+pyrimethamine triple therapy groups are significantly reduced, but loss of almost half of the group precludes practical application without further optimization (Fig. 7C). Interestingly, this experiment revealed that ELQ-334 combined with pyrimethamine impressively reduces brain cysts 95%, to near undetectable levels in C57BL/6 mice. This finding suggests that further optimization with ELQs paired with pyrimethamine or perhaps another anti-tachyzoite agent may be a promising avenue towards eradication of toxoplasmosis.

## Discussion

We recently showed that the efficacy of guanabenz against latent toxoplasmosis varies in different strains of mice. Guanabenz significantly reduces brain cysts in BALB/c mice, but in C57BL/6 mice, guanabenz treatment was lethal for half the group and showed an increase in cyst burden among the survivors (17). Here, we endeavored to prevent the loss of C57BL/6 mice through the co-administration of a second drug. The inclusion of pyrimethamine prevented the guanabenz-induced death; moreover, the C57BL/6 mice receiving the guanabenz+pyrimethamine co-treatment showed a significant reduction in brain cyst burden (Fig. 1C). This success encouraged us to pursue additional combination therapies with guanabenz in hopes of completely eliminating parasites in these mice.

Combination therapy is currently the standard of care against acute toxoplasmosis (7). First line drug combinations are pyrimethamine+sulfadiazine, which target two independent steps of the folic acid synthesis pathway, depriving the parasites of purines. While effective against acute toxoplasmosis, allergies or adverse effects to sulfa drugs force a switch to another therapeutic (23). The second line drug combination is pyrimethamine+clindamycin, the latter of which targets translation in the parasite-specific plastid-like organelle called the apicoplast (24).

Since pyrimethamine and clindamycin have different targets and are clinically available, we tested their efficacy with guanabenz against latent toxoplasmosis. Our results showed no improvement in brain cyst burden reduction in BALB/c mice. This was not surprising given that BALB/c mice, unlike C57BL/6 mice, showed no signs of reactivated toxoplasmosis or any indication that there were tachyzoites present during guanabenz treatment. Since both pyrimethamine and clindamycin rely on actively replicating parasites to be effective, the lack of tachyzoites in BALB/c mice likely explains the ineffectiveness of this combination therapy. It is possible that the residual population of cysts resides in an area of brain that is inaccessible to the drugs.

We also addressed the idea that co-administration of another cyst-reducing drug with guanabenz would have synergistic effects, potentially lowering cysts to a point below the threshold of detection. Interestingly, none of the compounds studied to date have been able to eradicate brain cysts completely as a monotherapy. Ours is the first study to examine a combination of two cyst-reducing drugs.

ELQs, which have a different mechanism of action than guanabenz, show a 76-88% reduction of brain cysts in CBA/J mice (14). *In vitro* studies showed a synergistic effect of guanabenz and ELQ-316 against tachyzoites as evidenced in two independent parasite growth assays (Fig. 3). Treatment of *in vitro* generated bradyzoite cysts resulted in gross morphological changes that were amplified when drugs were used in combination. Cysts treated with guanabenz were swollen with empty spaces and deformed bradyzoites visible within the cyst lumen, as previously reported (16). Cysts treated with ELQ-316 showed a shrunken morphology, resembling a raisin, with some parasites expressing SAG1. When the drugs were combined, there was an abundance of small, sometimes shrunken cysts that were expressing SAG1, in addition to larger shrunken cysts and a number of SAG1 positive parasites outside of cyst walls.

When we co-administered guanabenz and ELQ-334 (the prodrug of ELQ-316) *in vivo* against latent toxoplasmosis in BALB/c mice, we did not observe the additive effect we anticipated; in fact, we no longer observed the decrease in brain cyst burden normally seen in response to monotherapy with either drug. In other words, the drugs appear to be antagonistic *in vivo* through a mechanism that remains to be determined. The presence of SAG1 positive parasites seen *in vitro* with ELQ treatment could potentially explain its lack of effect when combined with guanabenz *in vivo*. We tested this possibility using a triple combination treatment by adding pyrimethamine, which we had previously shown to be efficacious in a system where tachyzoites are likely present as well. The triple drug combination was effective in significantly reducing cyst burden in BALB/c mice, but it did not lower it as much as the monotherapies.

Since we previously established that guanabenz works differently against latent toxoplasmosis in different mouse strains, we also performed combination drug treatments in C57BL/6 mice (17). It was unsurprising that guanabenz with ELQ-334 caused several of the mice to succumb to toxoplasmosis since this combination failed to reduce cysts in BALB/c mice. The addition of pyrimethamine did not help prevent the lethality seen in some of the mice given guanabenz+ELQ; however, the surviving mice displayed reduced cyst counts. Interestingly, the co-administration of pyrimethamine and ELQ-334 resulted in an impressive reduction in brain cyst burden in the C57BL/6 mice down towards undetectable limits.

In conclusion, while the addition of pyrimethamine to guanabenz treatment is effective in reducing cyst burden and preventing death in C57BL/6 mice, it is not more efficacious than guanabenz alone in BALB/c mice. The combination of guanabenz with ELQ-334, while showing additive effects *in vitro*, was not effective in either mouse model *in vivo*. However, the combination of ELQ-334 with pyrimethamine proved more effective than ELQ-334 alone, emerging as a promising combination that could be further optimized in the future.

## Acknowledgements

This research was supported by grants from the National Institutes of Health AI124723 and AI150616 (to W.J.S.) and the Career Development Award BX002440 and VA Merit Review Award BX004522 (to J.S.D.) J.M. was supported by PHS grant T32 AI060519 and the Joseph and Lucille Madri Family Scholarship.

